# Analysis of UK Biobank Cohort Reveals Novel Insights for Thoracic and Abdominal Aortic Aneurysms

**DOI:** 10.1101/2021.02.05.429911

**Authors:** Tamara Ashvetiya, Sherry X Fan, Yi-Ju Chen, Charles H Williams, Jeffery R. O’Connell, James A Perry, Charles C Hong

## Abstract

**Background:** Thoracic aortic aneurysm (TAA) and abdominal aortic aneurysm (AAA) are known to have a strong genetic component.

**Methods and Results:** In a genome-wide association study (GWAS) using the UK Biobank, we analyzed the genomes of 1,363 individuals with AAA compared to 27,260 age, ancestry, and sex-matched controls (1:20 case:control study design). A similar analysis was repeated for 435 individuals with TAA compared to 8,700 controls. Polymorphism with minor allele frequency (MAF) >0.5% were evaluated.

We identified novel loci near *LINC01021, ATOH8* and *JAK2* genes that achieved genome-wide significance for AAA (p-value <5×10^−8^), in addition to three known loci. For TAA, three novel loci in *CTNNA3, FRMD6* and *MBP* achieved genome-wide significance. There was no overlap in the genes associated with AAAs and TAAs. Additionally, we identified a linkage group of high-frequency variants (MAFs ∼10%) encompassing *FBN1*, the causal gene for Marfan syndrome, which was associated with TAA. In Finngen PheWeb, this *FBN1* haplotype was associated with aortic dissection. Finally, we found that baseline bradycardia was associated with TAA, but not AAA.

**Conclusions:** Our GWAS found that AAA and TAA were associated with distinct sets of genes, suggesting distinct underlying genetic architecture. We also found association between baseline bradycardia and TAA. These findings, including *JAK2* association, offer plausible mechanistic and therapeutic insights. We also found a common *FBN1* linkage group that is associated with TAA and aortic dissection in patients who do not have Marfan syndrome. These *FBN1* variants suggest shared pathophysiology between Marfan disease and sporadic TAA.

**Condensed Abstract:** In genome-wide association study (GWAS) of thoracic aortic aneurysm (TAA) and abdominal aortic aneurysm (AAA) using UK Biobank database, we found 3 novel loci associated with TAA, and 3 novel loci associated AAA. We also found significant association between baseline bradycardia and TAA. These findings, including *JAK2* association, offer plausible mechanistic and therapeutic insights. Additionally, we identified a common *FBN1* linkage group associated with TAA in patients who do not have Marfan syndrome. In the FinnGen cohort, this haplotype is associated with aortic dissection. These results suggest a shared pathophysiology between Marfan disease and sporadic TAA.

**Study Limitations:** As with any GWAS study, the discovery of novel loci associated with aortopathies does not prove functional causality, and the findings described herein needs to be validated by analysis of other databases, ideally in a patient population of more diverse genetic origins than the UK Biobank. The use of the ICD10 codes to classify disease carriers and noncarriers in a population cohort may not be the most accurate assessment of prevalence of aortopathies. The association between baseline bradycardia and TAA does not take into account the concurrent use of medications that may impact heart rate.

**Highlights:**

- Identification of 3 novel AAA-associated loci near *LINC01021, ATOH8* and *JAK2* genes.
- Identification of 3 novel TAA-associated loci near *CTNNA3, FRMD6* and *MBP* genes.
- Identification of a linkage group of common *FBN1* variants associated with non-syndromic TAA in the UK Biobank and with aortic dissection in the FinnGen cohort, strengthening the evidence for a shared pathophysiology between Marfan disease and nonsyndromic aortopathy.
- Association between baseline bradycardia and TAA but not AAA.

## Introduction

Aortic aneurysms (AA) carry a significant burden of morbidity and mortality. In 2018, thoracic aortic aneurysms (TAA) and abdominal aortic aneurysms (AAA) together were responsible for 9,923 deaths in the United States, typically from complications such as aortic dissection and rupture(1). AA primarily affect elderly males in the sixth and seventh decades of life with risk factors of tobacco use, hypertension and atherosclerosis(1). However, AAs found in patients younger than 65 years are more often attributed to genetic predisposition. For AAA, individuals with first-degree family members with the disease are at two-fold risk of developing AAA as compared to patients with no family history(2). Genetic studies of TAA and AAA have revealed genetic heterogeneity and polygenic inheritance patterns with variable disease penetrance(2–4).

For thoracic aortic aneurysms and dissection (TAAD), rare monogenetic syndromic disorders such as Marfan, Ehlers-Danlos and Loeys-Dietz syndromes are known to dramatically increase risk in younger individuals(5). This risk results from mutations in *FBN1, COL3A1*, and genes that encode TGF-β signaling proteins respectively(5). Beyond these, up to one-fifth of patients with TAAD have a familial predisposition toward aneurysmal disease(5). For instance, mutations in *FBN1* (fibrillin-1), the causal gene for Marfan syndrome, may increase the risk of thoracic aortic aneurysms or dissections even in individuals who do not have Marfan syndrome(4, 6, 7). Additionally, rare mutations in *ACTA2*, encoding smooth muscle protein alpha (α)-2 actin, may account for up to 14% of familial forms of TAAs(8). Furthermore, genetic studies of TAAD show it is closely associated with bicuspid aortic valve, another condition with strong heritability(5, 9).

Since AA has a strong genetic component in certain individuals, an enhanced understanding of these factors may ultimately aid the early detection of this silent disease before it progresses into life-threatening aortic dissections and ruptures. There is evidence to suggest that a strong personal or family history of aneurysms or dissection in individuals under the age of 50 should be an indication for genetic testing to diagnose inherited aortopathy(10). In appropriately selected patients with suspected familial aneurysmal disease, the yield of genetic testing could be as high as 36%(10). Yet, much of the underlying genetic risk factors remain unknown.

In this paper, we address the discovery of novel genetic loci that may portend increased risk for the development of thoracic and abdominal aortic aneurysms based on a genome-wide association study (GWAS) using data from the UK Biobank. The UK Biobank allows for powerful association studies with tremendous potential for expanding knowledge on the genetic basis of aortic aneurysms, as well as uncovering novel genetic variants associated with clinical diseases.

## Methods

### Ethical Approval

The present study, which involved deidentified data obtained from the UK Biobank Resource under Application Number 49852, received the proper ethical oversight, including the determination by the University of Maryland, Baltimore Institutional Review Board that the study is not human research (IRB #: HF-00088022).

### Study population

The UK Biobank, which was used for the GWAS presented here, is a large, ongoing prospective cohort study that recruited 502,682 UK participants between 2006-2010, ranging in age from 40-69 years at the time of recruitment. Extensive health-related records were collected from these participants, including clinical and genetic data, with over 820,000 genotyped single nucleotide polymorphisms (SNPs) and up to 90 million imputed variants available for most individuals. We carried out a genome-wide association study using the UK Biobank to interrogate the genome for statistically significant associations between SNPs and clinical manifestations of abdominal and thoracic aortic aneurysms at the population level.

### Genome-wide association study (GWAS)

Using data from the UK Biobank Resource on 487,310 subjects with imputed genotypes, we performed quality control by removing those with genetic relatedness exclusions (1,532), sex chromosome aneuploidy (651), mismatch between self-reported sex and genetically determined sex (372), recommended genomic analysis exclusions (480), and outliers for heterozygosity or missing rate (968). For “cases” we selected subjects with the following ICD10 (international classification of diseases) diagnostic codes, classified as either “main” or “secondary”: “abdominal aortic aneurysm, without mention of rupture” (1,363 patients, ICD10 code I71.4) and “abdominal aortic aneurysm, ruptured” (131 patients, ICD10 code I71.3). The selected set was purged of relatedness by removing one of each related pair in an iterative fashion until no related subjects remained. A pool of possible control subjects was generated by removing the cases and removing subjects with ICD10 codes listed as “excluded from controls” in **Supplementary Table 1**. Subjects with these ICD10 codes were removed from the population of controls to avoid introducing confounding factors, specifically the TAA and aortic valvular disorders, in the analysis. The pool was further reduced by removing related subjects. From the resulting reduced pool of possible controls, 20 control subjects were selected for each case subject, matched by sex, age and ancestry with sex as a required match (n=27,260 controls).

Incremental tolerances were used for age and ancestry with tolerances being expanded with each iteration until the desired number of matching controls were found for each case subject. The age tolerance ranged from 0 (exact match) up to 7 years. Ancestry matching was performed using principal components (PCs) supplied by the UK Biobank. The mathematical distance in a graph created by plotting PC1 x PC2 was used to test ancestry similarity. The ancestry “distance” tolerated ranged from 2 PC units up to a maximum of 80 PC units, where PC1 ranged from 0 to +400 and PC2 ranged from -300 to +100 units. Using these tolerances, 20 matching controls were found for every case. The analysis was repeated with patients who carried either a main or secondary diagnosis of “thoracic aortic aneurysm, without mention of rupture” (435 patients, ICD10 code I71.2) or “thoracic aortic aneurysm, ruptured” (22 patients, ICD10 code I71.1). Case to control ratio was again set at 1:20 (n=8,700 controls), and patients with the diagnoses listed in **Supplementary Table 2** were excluded from the control population. Subjects with these ICD10 codes were removed from the population of controls to avoid introducing confounding factors, specifically the AAA and aortic valvular disorders, in the analysis.

The association analysis was performed with PLINK2 using logistic regression(11). The thoracic aortic aneurysm (TAA) and abdominal aortic aneurysm (AAA) phenotypes were run against 40 million imputed variants supplied by the UK Biobank with imputation quality scores greater than 0.70. The analysis included covariates of sex, age, and principal components 1 through 5 to adjust for ancestry. Pre-calculated PC data was supplied by the UK Biobank. Our preliminary analysis showed that only the first 5 PCs had significance.

### Identification of significant SNPs for AAA and TAA Phenotypes

The SNPs identified in the analysis were filtered to include only those with minor allele frequency (MAF) of at least 0.5% and p-value <1 × 10^−6^, which is suggestive of genome-wide significance. The potential functional significance of associated variants was assessed by Eigen PC scores, presence of promoter or enhancer elements, and presence of DNase hypersensitivity sites in the affected regions of the genome. The Eigen PC score provides an aggregate score of functional importance for variants of interest, and is useful for prioritizing likely causal variants in a given genomic region. More positive Eigen PC scores are suggestive of more functional (disruptive) variants.

## Results

### Abdominal Aortic Aneurysm

Of the 1,363 individuals documented to have abdominal aortic aneurysms, 131 (9.61%) had rupture of the aneurysm (**Table 1**). The affected patients ranged in age at diagnosis from 42.42 to 79.04 years (mean = 68.08); 86.72% were male and 13.28% female. Genetic ancestry was predominantly British (93.54%); however, patients were also represented from Irish (2.64%), Indian (0.22%), Caribbean (0.51%), and African (0.15%) backgrounds. Based on GWAS of these 1,363 individuals, we identified four independent loci near the *LINC01021, ADAMTS8, ATOH8* and *JAK2* genes (7 SNPs in total) that achieved a genome-wide significance level of p-value <5 × 10^−8^ for variants with MAF ≥0.5% (**Figure 1A**). Of these, *LINC01021, ADAMTS8* and *JAK2* represent novel loci associated with AAA, and each of the four loci harbors SNPs that possess features suggestive of functional importance, with biological plausibility as disease susceptibility loci (**Table 2**). In addition, we found several distinct variants that do not reach the definitive threshold for genome-wide significance but nevertheless possess a strong basis for biologic plausibility and are within the suggestive threshold for genome-wide significance (p-value <1e^−6^; **Table 2**). Full GWAS results for AAA are included in **Supplementary Table 5** for variants with MAF ≥0.5% and p-value <1 × 10^−6^.

**Table 1.** Basic demographics of individuals with abdominal aortic aneurysm (AAA) and thoracic aortic aneurysm (TAA) in UK Biobank.

|  | **Individuals**<br>**(#)** | **# with**<br>**rupture** | **Mean Age at**<br>**Diagnosis**<br>**(yr)** | **Male (%)** | **British**<br>**Ancestry (%)** |
| --- | --- | --- | --- | --- | --- |
| **AAA** | 1363 | 131 (9.6%) | 68.1 | 86.7 | 93.5 |
| **TAA** | 435 | 22 (5.1%) | 65.3 | 68.7 | 87.6 |

**Figure 1.**
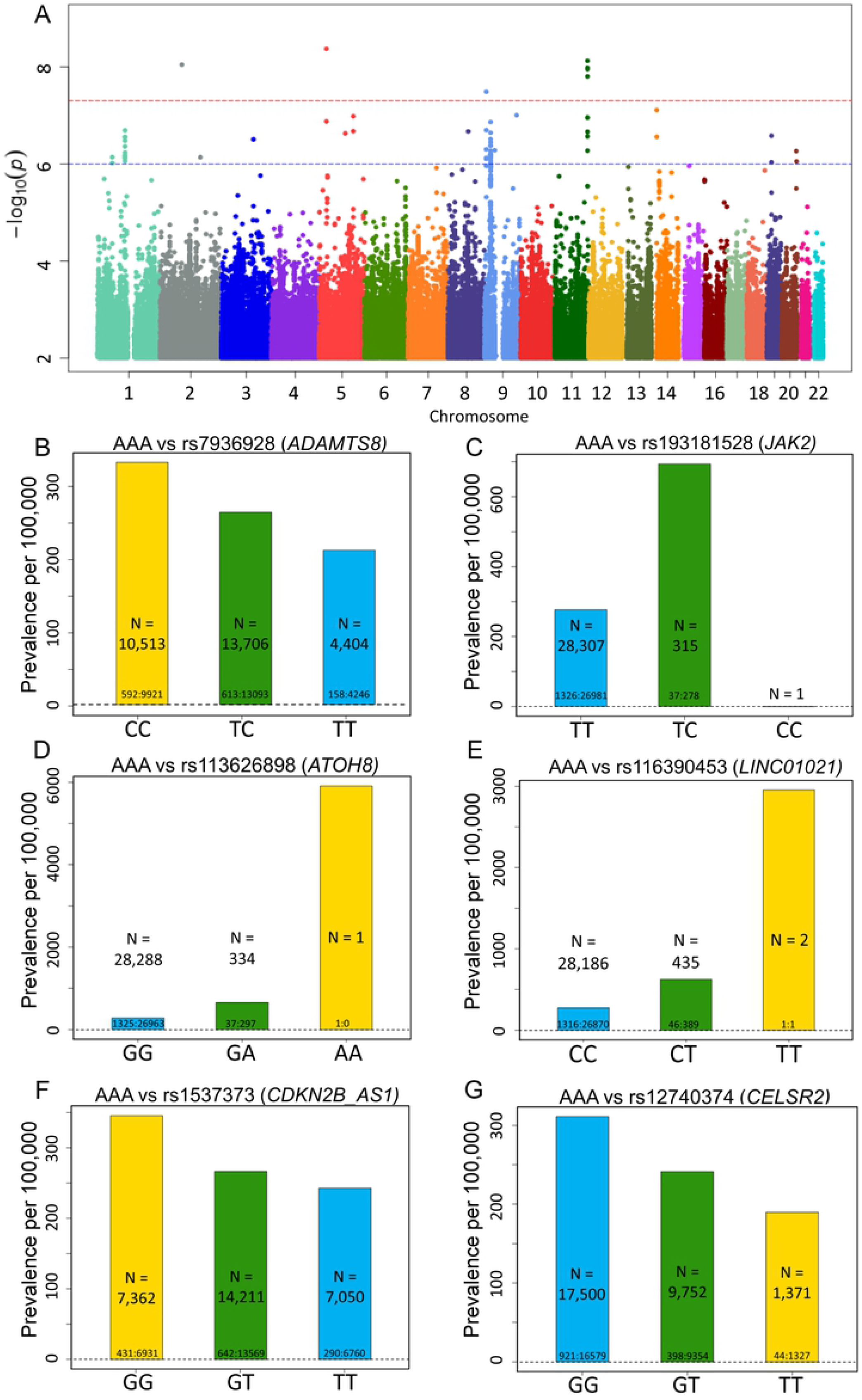
Top SNPs associated with AAA. (**A**) Manhattan plot of GWAS results (MAF >0.5%) for AAA. Significance is displayed on the y-axis as -log_10_ of the p-value, with results ordered along the x-axis by chromosome (each bar represents a different chromosome). **(B-G)** Prevalence of abdominal aortic aneurism (AAA) per 100,000 participants in the UK Biobank by genotype. Bars labeled with ratio of cases: controls. (**B**) Prevalence of AAA decreases with *ADAMTS8* variant rs7936928 status (P-value = 7.51×10^− 9^, OR per T allele = 0.786). Decrease in AAA prevalence is noted in the homozygotes for the minor allele (T/T) in comparison to the heterozygotes (C/T) and the noncarriers (C/C) in a stepwise, “dosage-dependent” manner. **(C)** Prevalence of AAA increases with *JAK2* variant rs193181528 status (P-value = 3.26×10^−8^, OR per C allele = 2.776). **(D)** Prevalence of AAA increases with *ATOH8* variant rs113626898 status (P-value = 9.06×10^−9^, OR per A allele = 2.714). **(E)** Prevalence of AAA increases with *LINC01021* variant rs116390453 status (P-value = 4.26 ×10^−9^, OR per T allele = 2.505). **(F)** Prevalence of AAA decreases with *CDKN2B-AS1* variant rs1537373 status (P-value = 6.68×10^−7^, OR per T allele = 0.8211). **(G)** Prevalence of AAA decreases with *CELSR2* variant rs12740374 status (P-value = 2.04×10^−7^, OR per T allele = 0.7668).

**Table 2.**
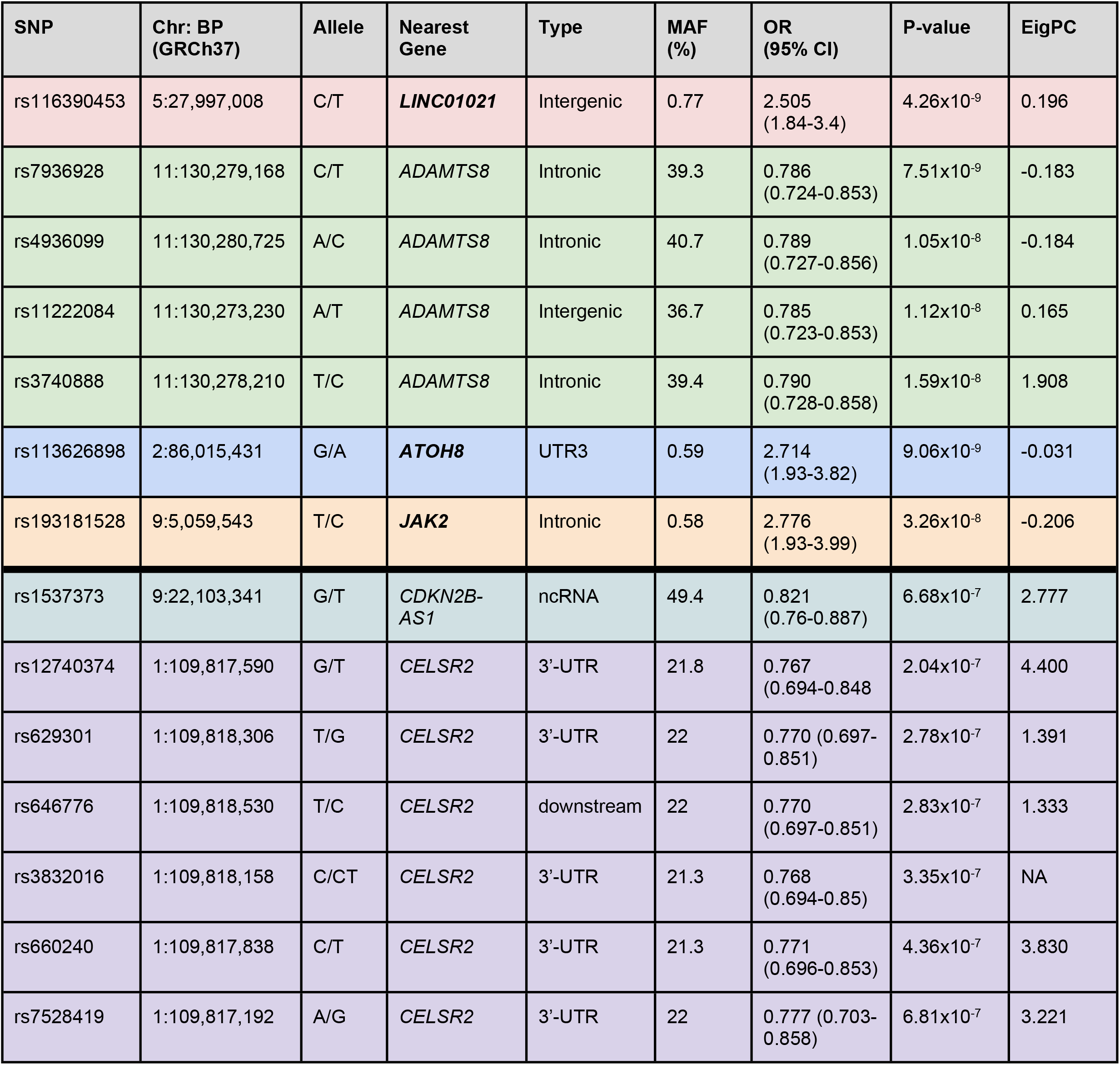
Top SNPs associated with abdominal aortic aneurysm. 7 variants in 4 genes, *LINC01021, ADAMTS8, ATOH8* and J*AK2*, reached genome-wide significance P-value of < 5 ×10^−8^, while 7 additional variants in *CDKN2B-AS1* and *CELSR2*, while not statistically significant, replicated findings from earlier studies. Of these, *LINC01021, ATOH8* and J*AK2* are novel AAA-associated loci identified in the present study (bold faced). Chr:BP denotes the chromosome location and NCBI Build 37 SNP physical position. Variants that are in linkage disequilibrium (LD) are identically colored. MAF, minor allele frequency. OR, odds ratio. EigPC, Eigen PC score, which provides an aggregate score of functional importance for variants of interest; more positive Eigen PC scores are suggestive of more functional (disruptive) variants.

| SNP | Chr: BP (GRCh37) | Allele | Nearest Gene | Type | MAF (%) | OR (95% CI) | P-value | EigPC |
| --- | --- | --- | --- | --- | --- | --- | --- | --- |
| rs116390453 | 5:27,997,008 | C/T | **LINC01021** | Intergenic | 0.77 | 2.505 (1.84-3.4) | 4.26x10<sup>-9</sup> | 0.196 |
| rs7936928 | 11:130,279,168 | C/T | ADAMTS8 | Intronic | 39.3 | 0.786 (0.724-0.853) | 7.51x10<sup>-9</sup> | -0.183 |
| rs4936099 | 11:130,280,725 | A/C | ADAMTS8 | Intronic | 40.7 | 0.789 (0.727-0.856) | 1.05x10<sup>-8</sup> | -0.184 |
| rs11222084 | 11:130,273,230 | A/T | ADAMTS8 | Intergenic | 36.7 | 0.785 (0.723-0.853) | 1.12x10<sup>-8</sup> | 0.165 |
| rs3740888 | 11:130,278,210 | T/C | ADAMTS8 | Intronic | 39.4 | 0.790 (0.728-0.858) | 1.59x10<sup>-8</sup> | 1.908 |
| rs113626898 | 2:86,015,431 | G/A | **ATOH8** | UTR3 | 0.59 | 2.714 (1.93-3.82) | 9.06x10<sup>-9</sup> | -0.031 |
| rs193181528 | 9:5,059,543 | T/C | **JAK2** | Intronic | 0.58 | 2.776 (1.93-3.99) | 3.26x10<sup>-8</sup> | -0.206 |
| rs1537373 | 9:22,103,341 | G/T | CDKN2B-AS1 | ncRNA | 49.4 | 0.821 (0.76-0.887) | 6.68x10<sup>-7</sup> | 2.777 |
| rs12740374 | 1:109,817,590 | G/T | CELSR2 | 3'-UTR | 21.8 | 0.767 (0.694-0.848) | 2.04x10<sup>-7</sup> | 4.400 |
| rs629301 | 1:109,818,306 | T/G | CELSR2 | 3'-UTR | 22 | 0.770 (0.697-0.851) | 2.78x10<sup>-7</sup> | 1.391 |
| rs646776 | 1:109,818,530 | T/C | CELSR2 | downstream | 22 | 0.770 (0.697-0.851) | 2.83x10<sup>-7</sup> | 1.333 |
| rs3832016 | 1:109,818,158 | C/CT | CELSR2 | 3'-UTR | 21.3 | 0.768 (0.694-0.85) | 3.35x10<sup>-7</sup> | NA |
| rs660240 | 1:109,817,838 | C/T | CELSR2 | 3'-UTR | 21.3 | 0.771 (0.696-0.853) | 4.36x10<sup>-7</sup> | 3.830 |
| rs7528419 | 1:109,817,192 | A/G | CELSR2 | 3'-UTR | 22 | 0.777 (0.703-0.858) | 6.81x10<sup>-7</sup> | 3.221 |

Among the SNPs with genome-wide significance, we identified a linkage group of 4 variants in close proximity to the ADAM metallopeptidase with thrombospondin type 1 motif 8 gene (*ADAMTS8*), which encodes an inflammation-regulated enzyme expressed in macrophage-rich areas of atherosclerotic plaques(12). Prior studies have described upregulation of *ADAMTS8* in the macrophages of patients with abdominal aortic aneurysms(13). The significant linkage group of *ADAMTS8* variants that we identified includes rs7936928 (intronic), rs4936099 (intronic), rs11222084 (intergenic), and rs3740888 (intronic); these variants have p-values 7.51 × 10^−9^-1.59 × 10^−8^, MAF 36.7%-40.7%, and odds ratios 0.785-0.790 (**Table 2; Figure 1B**). Of note, the *ADAMTS8* locus was recently identified among 14 novel AAA-risk loci identified from a study of the Million Veteran Program(14).

Other notable variants identified in this genome-wide analysis include the intronic variant rs193181528 (**Table 2**; **Figure 1C**; p-value 3.26 × 10^−8^, MAF 0.58%, OR 2.776), located within the gene encoding JAK2 tyrosine kinase. Mutations in *JAK2* are suspected to play a potential role in the progression of AAAs(15, 16). Among human aortic tissues collected from patients undergoing AAA surgery, *JAK2* expression levels were higher in patients with AAA as compared to controls(15). Treatment with JAK2/STAT3 pathway inhibitors attenuated experimental AAA progression by reducing the expression of pro-inflammatory cytokines and matrix metalloproteinases as well as inflammatory cell infiltration(15, 16).

The analysis also identified variant rs113626898 (**Table 2**; **Figure 1D**; p-value 9.06 ×10^− 9^, MAF 0.59%, OR 2.714) in the gene encoding atonal bHLH transcription factor 8 (*ATOH8*), which plays a role in myogenesis and contributes to endothelial cell differentiation, proliferation and migration(17). Dysregulation of this gene could plausibly contribute to aneurysmal formation given its important role in the cell cycles of myocytes and endothelial cells.

Variant rs116390453 (**Table 2; Figure 1E**; p-value 4.26 ×10^−9^, MAF 0.77%, OR 2.505) is within a long intergenic non-coding RNA (*LINC01021*), also known as p53 upregulated regulator of p53 levels (PURPL). To date, PURPL has not been linked to aneurysmal formation.

In addition, we identified several distinct variants that do not reach the standard threshold for genome-wide significance for association with AAA, but are nevertheless within the suggestive threshold for genome-wide significance (p-value <1 × 10^−6^) and possess a strong basis for biologic plausibility (**Table 2**). Variant rs1537373 (**Figure 1F**; p-value 6.68 × 10^−7^, MAF 49.9%, OR 0.821) in *CDKN2B-AS1*, encoding the long non-coding RNA known as cyclin dependent kinase inhibitor CDKN2B antisense RNA1, is located within the *CDKN2B-CDKN2A* gene cluster at chromosome 9p21, a major genetic susceptibility locus for coronary artery disease, atherosclerosis, and myocardial infarction(18). This locus has also been previously associated with intracranial aneurysm and AAA formation(14, 19–22). Thus, *CDKN2B-AS1* variant rs1537373 may increase the risk of AAA formation indirectly through the development of atherosclerosis, a major clinical risk factor for AAA.

Finally, our analysis identified a linkage group of six SNPs within the *CELSR2* gene, encoding the cadherin EGF LAG seven-pass G-type receptor 2 (**Table 2**; **Figure 1G**; p-values 2.04 ×10^−7^-6.81 ×10^−7^, MAF 21.3-22%, OR 0.767-0.777). While *CELSR2* SNPs do not meet traditional P-value cutoff for genome-wide significance, our findings corroborate prior GWAS associations of *CELSR2* with AAA(14, 23, 24), and identify new common SNPs that are associated with AAA (specifically rs3832016 and rs660240). Of these variants, rs12740374, rs660240 and rs7528419 have a particularly high Eigen PC score of 4.4, 3.8 and 3.2, respectively, suggesting a functional role (**Table 2**).

### Thoracic Aortic Aneurysm

Of the 435 individuals with thoracic aortic aneurysms, 22 (5.06%) had rupture of the aneurysm (**Table 1**). The affected patients ranged in age at diagnosis from 36.47 to 78.65 years (mean = 65.28); 68.74% were male and 31.26% female, with genetic ancestry of British (87.59%), Irish (2.07%), Indian (1.61%), Caribbean (1.38%), and African (0.23%) origins. Based on GWAS, we identified three SNPs that achieved a genome-wide significance level (p-value <5 × 10^−8^) together with a MAF ≥0.5% (**Figure 2A; Table 3**). Full GWAS results for TAA are included in **Supplementary Table 6** for variants with MAF ≥ 0.5%.

**Figure 2.**
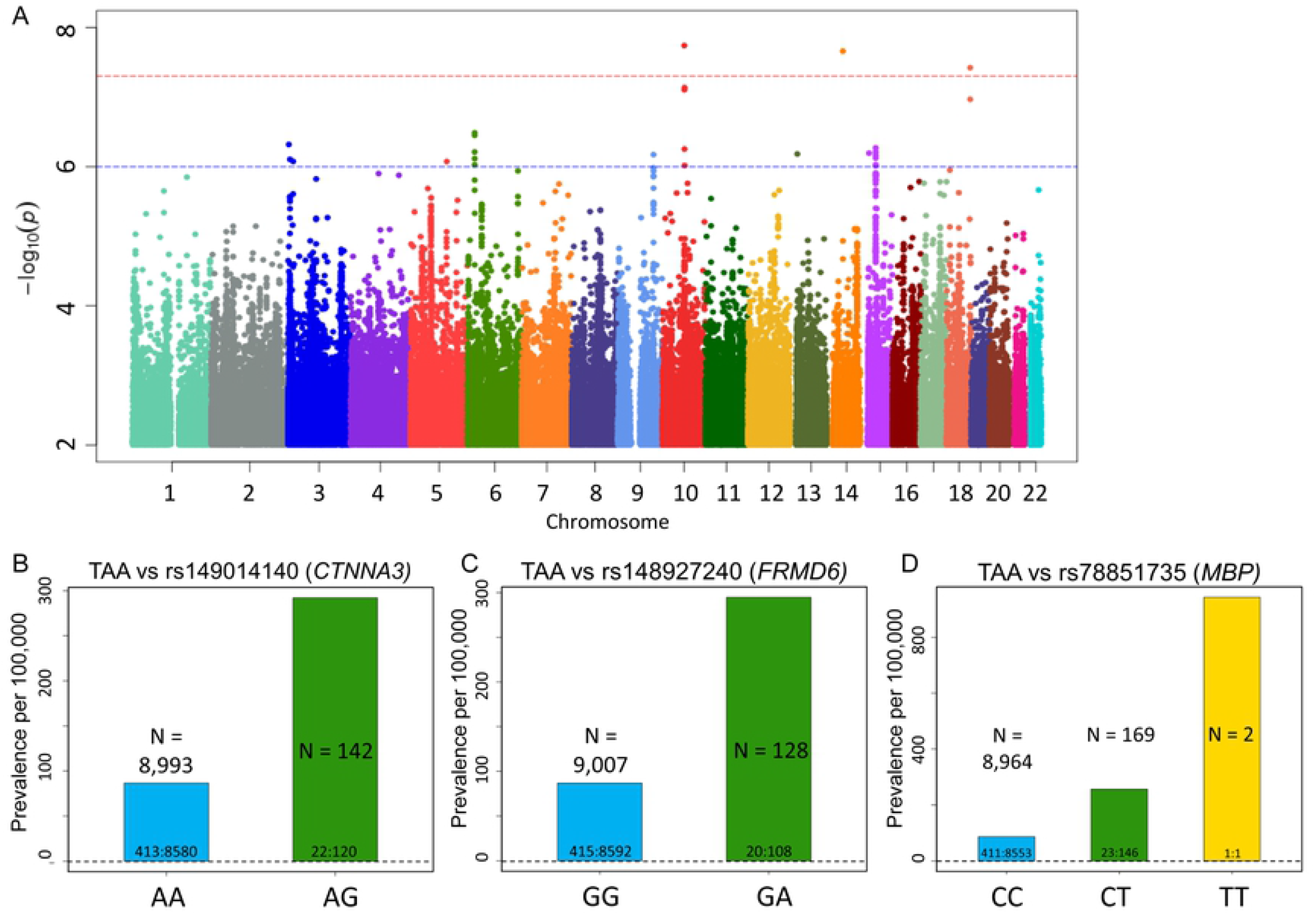
Top SNPs associated with TAA. **(A)** Manhattan plot of GWAS results (MAF >0.5%) for TAA. **(B-D)** Prevalence of thoracic aortic aneurism (AAA) per 100,000 participants in the UK Biobank by genotype. Bars labeled with ratio of cases: controls. **(B)** Prevalence of TAA increases with *CTNNA3* variant rs149014140 status (P-value = 1.82×10^−8^, OR per G allele = 4.268). **(C)** Prevalence of TAA increases with *FRMD6* variant rs148927240 status (P-value = 2.19×10^−8^, OR per A allele = 4.23). **(D)** Prevalence of TAA increases with *MPB* variant rs78851735 status (P-value = 3.79×10^−8^, OR per T allele = 3.446).

**Table 3.**
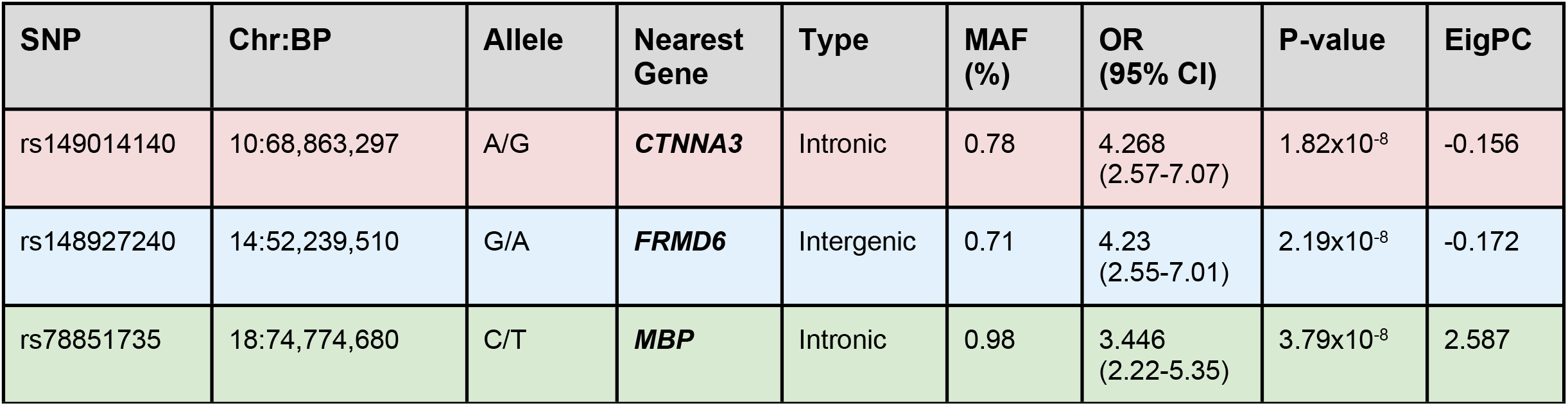
SNPs in 3 novel loci associated with thoracic aortic aneurysm. *CTNNA3, FRMD6* and *MBP* are novel TAA-associated loci identified in the present study (bold faced). Chr:BP denotes the chromosome location and NCBI Build 37 SNP physical position. MAF, minor allele frequency; OR, odds ratio. EigPC, Eigen PC score.

| SNP | Chr:BP | Allele | Nearest Gene | Type | MAF (%) | OR (95% CI) | P-value | EigPC |
| --- | --- | --- | --- | --- | --- | --- | --- | --- |
| rs149014140 | 10:68,863,297 | A/G | **CTNNA3** | Intronic | 0.78 | 4.268<br>(2.57-7.07) | 1.82x10<sup>-8</sup> | -0.156 |
| rs148927240 | 14:52,239,510 | G/A | **FRMD6** | Intergenic | 0.71 | 4.23<br>(2.55-7.01) | 2.19x10<sup>-8</sup> | -0.172 |
| rs78851735 | 18:74,774,680 | C/T | **MBP** | Intronic | 0.98 | 3.446<br>(2.22-5.35) | 3.79x10<sup>-8</sup> | 2.587 |

Among the SNPs with significant p-values, variant rs149014140 in *CTNNA3* gene is of particular interest (**Figure 2B**; p-value 1.82 × 10^−8^, MAF 0.78%, OR 4.268) since *CTNNA3* encodes a vinculin/alpha-catenin family protein known to play a role in cell-to-cell adhesion of muscle cells(25). Another significant variant is rs148927240 (**Figure 2C**; p-value 2.19 × 10^−8^, MAF 0.71%, OR 4.23), an intergenic variant located between the long non-coding RNA FERM domain containing 6 (*FRMD6*), involved in cell contact inhibition and cell cycle regulation(26), and the gene encoding one of the gamma subunits of a guanine nucleotide-binding protein (*GNG2*). Finally, variant rs78851735 (**Figure 2D**; p-value 3.79 × 10^−8^, MAF 0.98%, OR 3.446) is an intronic variant within the gene encoding myelin basic protein (*MBP*), which is the major protein in myelin sheaths of the nervous system(27). The biological relevance of *FRMD6, GNG2* and *MBP* with respect to the development of aortic aneurysms is unclear at this time.

In addition, we identified we identified a linkage group of high-frequency variants (MAFs 9.56-9.97%, odds ratios 1.615-1.644) that do not reach the standard threshold for genome-wide significance for association with TAA, but nevertheless have a strong basis for biologic plausibility since they fall in a large region of linkage disequilibrium encompassing *FBN1* (**Table 5**). *FBN1* encodes the fibrillin-1 protein and is implicated in the pathogenesis of Marfan syndrome as well as other disorders of the connective tissues including ectopia lentis syndrome, Weill-Marchesani syndrome, Shprintzen-Goldberg syndrome and neonatal progeroid syndrome(28). Fibrillin-1 is important in maintaining the integrity of connective tissues throughout the body, as it serves as a structural component of calcium-binding myofibrils(28). Intronic variant rs1561207 (**Figure 3A**; p-value 1.57 ×10^−6^, MAF 9.89%, OR 1.615) is of special interest given its high Eigen PC score. This variant is in near-perfect linkage with the seven other *FBN1* variants, all of which meet the suggestive statistical threshold for genome-wide significance (p-value <1 × 10^−6^) (**Table 4**).

**Figure 3.**
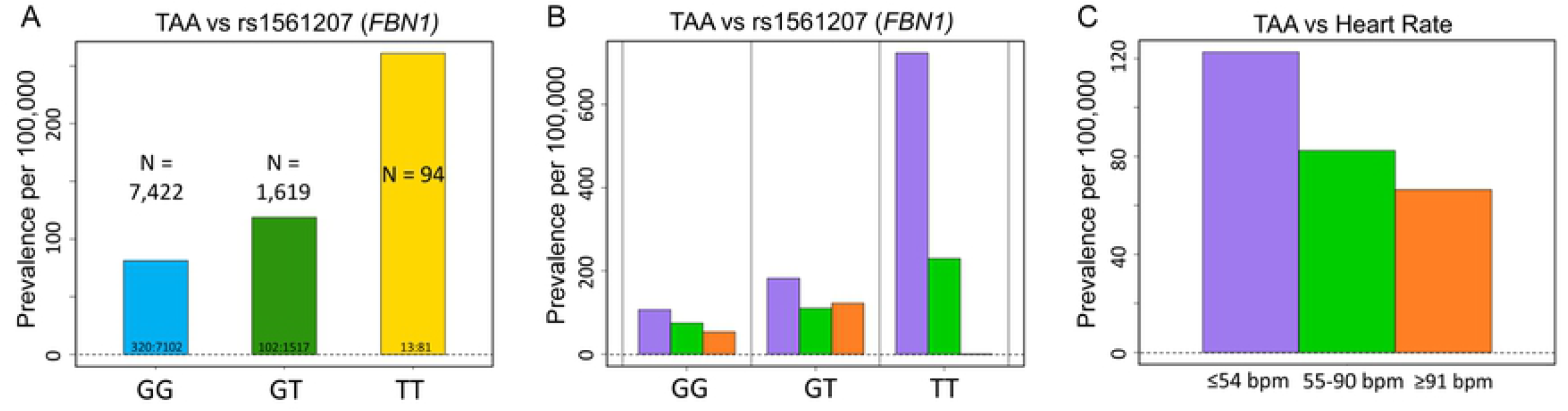
Prevalence of TAA increases with *FBN1* variant rs1561207 status and bradycardia. **(A)** Increase in TAA prevalence per 100,000 participants in the UK Biobank is noted in the homozygotes for the minor allele (T/T) in comparison to the heterozygotes (C/T) and the noncarriers (C/C) in a pronounced stepwise, “dosage-dependent” manner (P-value = 1.57×10^−6^, OR per T allele 1.615). Bars labeled with ratio of cases: controls. **(B)** Prevalence of TAA rises with bradycardia (purple, heart rate ≤ 54 beats per minute, bpm) in both *FBN1* variant carriers and noncarriers, but the impact of bradycardia is more dramatic in homozygous variant carriers. **(C)** Irrespective of genotype, TAA prevalence appears to increase in a stepwise manner from tachycardia (orange, defined as heart rate ≥ 91 bpm, OR = 1.89) to normal rate (green, heart rate 55 to 90 bpm, OR = 1.62) to bradycardia (purple, heart rate ≤ 54 bpm, OR = 2.09). P-value = 0.01622 by Pearson’s Chi-squared test.

**Table 4.** Linkage group of *FBN1* variants associated with thoracic aortic aneurysm. Chr:BP denotes the chromosome location and NCBI Build 37 SNP physical position. MAF, minor allele frequency; OR, odds ratio. EigPC, Eigen PC score.

| SNP | Chr: BP | Allele | Type | MAF (%) | OR (95% CI) | P-value | EigPC |
| --- | --- | --- | --- | --- | --- | --- | --- |
| rs1561207 | 15: 48,858,971 | G/T | Intronic | 9.89 | 1.615 (1.33-1.96) | 1.57x10<sup>-6</sup> | 0.495 |
| rs689304 | 15: 48,922,360 | C/T | Intronic | 9.93 | 1.642 (1.35-1.99) | 5.38x10<sup>-7</sup> | -0.037 |
| rs625034 | 15: 48,926,202 | T/C | Intronic | 9.9 | 1.64 (1.35-1.99) | 6.15x10<sup>-7</sup> | -0.184 |
| rs1036476 | 15: 48,914,775 | T/C | Intronic | 9.89 | 1.636 (1.35-1.99) | 6.93x10<sup>-7</sup> | -0.126 |
| rs2028109 | 15: 48,919,103 | A/C | Intronic | 9.9 | 1.634 (1.35-1.99) | 7.54x10<sup>-7</sup> | 0.013 |
| rs2455925 | 15: 48,893,649 | T/C | Intronic | 9.97 | 1.625 (1.34-1.97) | 9.37x10<sup>-7</sup> | -0.188 |
| rs4775769 | 15: 48,939,888 | G/T | Intergenic | 9.56 | 1.644 (1.35-2.01) | 9.70x10<sup>-7</sup> | -0.229 |

Interestingly, each of these *FBN1* variants demonstrated a pronounced dose-dependence: homozygotes had significantly higher prevalence of thoracic aortic aneurysm than heterozygotes (**Figure 3A**). Comorbid conditions of TAA patients with these *FBN1* variants (**Supplementary Table 3**), as well as gender and age at diagnosis (**Supplementary Table 4**) were similar to the overall population of TAA patients. In FinnGen PheWeb, comprised of genetic and clinical information on 178,899 Finnish cohort, this haplotype was associated with aortic dissection (p =2.3 × 10^−5^; **Supplementary Figure 1**). Interestingly, GWAS of thoracic aorta images in the UK Biobank revealed similar association of FBN1 with dilated aorta(29). Thus, our study strengthens the emerging functional association between *FBN1* and nonsyndromic aortopathy(30).

### Relationship between pulse rate and prevalence of aortic aneurysms

When we analyzed AA prevalence by baseline pulse rate for each of the statistically significant SNPs identified in this study, an unexpected correlation emerged between heart rate and prevalence of aortic aneurysms. For TAA, but not AAA, there was a general trend toward increased prevalence in individuals with bradycardia (defined as heart rate ≤54 beats per minute), regardless of genotype (**Figure 3C**). Interestingly, this trend seems more marked for homozygous carriers of *FBN1* variants (**Figure 3B**). In contrast, a general trend toward slightly increased AAA prevalence is seen with tachycardia, although the effect is smaller overall (**Supplementary Figure 2**).

## Discussion

Genome wide association analysis of abdominal and thoracic aortic aneurysmal disease in the UK Biobank revealed novel loci near *LINC01021, ATOH8* and *JAK2* genes associated with AAA, and novel loci near *CTNNA3, FRMD6* and *MBP* genes associated with TAA. Out of the 24 loci previously established for AAA, three were replicated by our analysis, *ADAMTS8, CELSR2* and *CDKN2B-AS1*(14, 31). Based on the data compiled here with the thresholds for p-value and minor allele frequency as set forth in the methods section, there was no significant overlap in the SNPs associated with AAAs and those associated with TAAs. This suggests a distinct underlying genetic architecture, and a distinct pathophysiology, of these two aortopathies.

A potentially clinically relevant finding is the identification of a linkage group of SNPs encompassing the *FBN1* gene, which is associated with TAA in the UK Biobank and with aortic dissection in FinnGen cohorts. This haplotype demonstrated a pronounced dose-dependence, with homozygous carriers associated with ∼2.2-fold higher prevalence of thoracic aortic aneurysm than heterozygotes. Given the relatively high minor allele frequencies for these SNPs (9.56-9.97%), as well as the well-defined role of *FBN1* in the pathogenesis of connective tissues disorders including Marfan syndrome, we hypothesize that mutations within this linkage group may account for a non-trivial portion of nonsyndromic thoracic aortic aneurysms and dissections, particularly those within the context of positive family history. Therefore, these variants could merit inclusion in genetic screening panels for familial thoracic aneurysmal disease. Taken together with the findings of Pirruccello, et al., our finding suggests some degree of shared pathophysiology between aortic disease in Marfan syndrome and sporadic thoracic aortic aneurysm(29).

An unexpected finding of this study is the apparent association of TAA prevalence and baseline bradycardia. It is unknown whether bradycardia is a consequence of beta-adrenergic blocker usage in those diagnosed with TAA. Nevertheless, the fact that this association is seen only with TAA, but not AAA, suggests some biological basis. For example, widening of pulse pressure accompanying bradycardia may disproportionately increase wall tension in the proximal thoracic aorta but not the distal abdominal aorta. The possibility that bradycardia somehow contributes to TAA formation and therefore should be avoided in susceptible individuals, particularly with *FBN* variants, may warrant further investigation.

The approach used in this paper has several limitations. As with any GWAS study, the discovery of novel loci associated with aortopathies does not prove functional causality, and the findings described herein needs to be validated by analysis of other databases, ideally in a patient population of more diverse genetic origins than the UK Biobank. Nonetheless, some of these loci, such *JAK2*, suggest plausible mechanistic and therapeutic insights that merit further investigation. We also note that our study identified fewer associations than recent GWAS studies on the Million Veterans Program (MVP) cohort and on aortic images in the UK Biobank(14, 29). The fewer associations we identified compared to the MVP study may due to the fact that we examined 1,363 patients with AAA whereas the MVP study examined 7,642 patients(14). Additionally, the greater number of associations found by Pirruccello et al. could reflect the fact that they used imaging to identify individuals with subclinical aortopathy not captured by ICD10 codes(29). Finally, we also note that, in addition to the variants and loci discussed here, there are many more that didn’t make the genome-wide significance cutoff of p < 5 × 10^−8^ or MAF cutoff ≥ 0.5%. Thus, much of the genetic underpinnings of abdominal and thoracic aortic aneurysm formation remain to be discovered.

## Acknowledgements

This research was conducted using the UK Biobank Resource under Application Number 49852. This work was supported by NIGMS R01GM118557 and NHLBI R01HL1351291 to CCH, and NHLBI 1U01HL137181 to JP. The funders had no role in the design and conduct of the study; collection, management, analysis and interpretation of the data; preparation, review or approval of the manuscript; or decision to submit the manuscript for publication.

## Abbreviations

AA: (aortic aneurysm)
AAA: (abdominal aortic aneurysm)
GWAS: (genome-wide association study)
ICD: (international classification of diseases)
LD: (linkage disequilibrium)
MAF: (minor allele frequency)
MVP: (Million Veteran Program)
PC: (principal component)
SNP: (single nucleotide polymorphism)
TAA: (thoracic aortic aneurysm)
TAAD: (thoracic aortic aneurysms and dissection)
UK: (United Kingdom)

## References

1. CDC. Aortic Aneurysm | cdc.gov. Centers for Disease Control and Prevention 2020. Available at :https://www.cdc.gov/heartdisease/aortic_aneurysm.htm. Accessed October 3, 2020.

2. Shibamura H, Olson JM, van Vlijmen-Van Keulen C, et al. Genome scan for familial abdominal aortic aneurysm using sex and family history as covariates suggests genetic heterogeneity and identifies linkage to chromosome 19q13. Circulation 2004;109:2103–2108.

3. Guo D, Hasham S, Kuang SQ, et al. Familial thoracic aortic aneurysms and dissections: genetic heterogeneity with a major locus mapping to 5q13-14. Circulation 2001;103:2461– 2468.

4. Vaughan CJ, Casey M, He J, et al. Identification of a chromosome 11q23.2-q24 locus for familial aortic aneurysm disease, a genetically heterogeneous disorder. Circulation 2001;103:2469–2475.

5. Ostberg NP, Zafar MA, Ziganshin BA, Elefteriades JA. The Genetics of Thoracic Aortic Aneurysms and Dissection: A Clinical Perspective. Biomolecules 2020;10.

6. Milewicz DM, Michael K, Fisher N, Coselli JS, Markello T, Biddinger A. Fibrillin-1 (FBN1) mutations in patients with thoracic aortic aneurysms. Circulation 1996;94:2708–2711.

7. LeMaire SA, McDonald M-LN, Guo D-C, et al. Genome-wide association study identifies a susceptibility locus for thoracic aortic aneurysms and aortic dissections spanning FBN1 at 15q21.1. Nat Genet 2011;43:996–1000.

8. Guo D-C, Pannu H, Tran-Fadulu V, et al. Mutations in smooth muscle alpha-actin (ACTA2) lead to thoracic aortic aneurysms and dissections. Nat Genet 2007;39:1488–1493.

9. Gould RA, Aziz H, Woods CE, et al. ROBO4 variants predispose individuals to bicuspid aortic valve and thoracic aortic aneurysm. Nat Genet 2019;51:42–50.

10. Hicks KL, Byers PH, Quiroga E, Pepin MG, Shalhub S. Testing patterns for genetically triggered aortic and arterial aneurysms and dissections at an academic center. J Vasc Surg 2018;68:701–711.

11. Chang CC, Chow CC, Tellier LC, Vattikuti S, Purcell SM, Lee JJ. Second-generation PLINK: rising to the challenge of larger and richer datasets. Gigascience 2015;4:7.

12. Wågsäter D, Björk H, Zhu C, et al. ADAMTS-4 and -8 are inflammatory regulated enzymes expressed in macrophage-rich areas of human atherosclerotic plaques. Atherosclerosis 2008;196:514–522.

13. Lamblin N, Ratajczak P, Hot D, et al. Profile of macrophages in human abdominal aortic aneurysms: a transcriptomic, proteomic, and antibody protein array study. J Proteome Res 2010;9:3720–3729.

14. Klarin D, Verma SS, Judy R, et al. Genetic Architecture of Abdominal Aortic Aneurysm in the Million Veteran Program. Circulation 2020;142:1633–1646.

15. Xiao J, Wei Z, Chen X, et al. Experimental abdominal aortic aneurysm growth is inhibited by blocking the JAK2/STAT3 pathway. Int J Cardiol 2020;312:100–106.

16. Wu Q-Y, Cheng Z, Zhou Y-Z, et al. A novel STAT3 inhibitor attenuates angiotensin II-induced abdominal aortic aneurysm progression in mice through modulating vascular inflammation and autophagy. Cell Death Dis 2020;11:131.

17. Fang F, Wasserman SM, Torres-Vazquez J, et al. The role of Hath6, a newly identified shear-stress-responsive transcription factor, in endothelial cell differentiation and function. J Cell Sci 2014;127:1428–1440.

18. Roberts R, Stewart AFR. 9p21 and the genetic revolution for coronary artery disease. Clin Chem 2012;58:104–112.

19. Samuel N, Radovanovic I. Genetic basis of intracranial aneurysm formation and rupture: clinical implications in the postgenomic era. Neurosurg Focus 2019;47:E10.

20. Hitchcock E, Gibson WT. A Review of the Genetics of Intracranial Berry Aneurysms and Implications for Genetic Counseling. J Genet Couns 2017;26:21–31.

21. Helgadottir A, Thorleifsson G, Manolescu A, et al. A common variant on chromosome 9p21 affects the risk of myocardial infarction. Science 2007;316:1491–1493.

22. Helgadottir A, Thorleifsson G, Magnusson KP, et al. The same sequence variant on 9p21 associates with myocardial infarction, abdominal aortic aneurysm and intracranial aneurysm. Nat Genet 2008;40:217–224.

23. Jones GT, Bown MJ, Gretarsdottir S, et al. A sequence variant associated with sortilin-1 (SORT1) on 1p13.3 is independently associated with abdominal aortic aneurysm. Hum Mol Genet 2013;22:2941–2947.

24. Carino D, Sarac TP, Ziganshin BA, Elefteriades JA. Abdominal Aortic Aneurysm: Evolving Controversies and Uncertainties. Int J Angiol 2018;27:58–80.

25. van Hengel J, Calore M, Bauce B, et al. Mutations in the area composita protein αT-catenin are associated with arrhythmogenic right ventricular cardiomyopathy. Eur Heart J 2013;34:201– 210.

26. Xu Y, Wang K, Yu Q. FRMD6 inhibits human glioblastoma growth and progression by negatively regulating activity of receptor tyrosine kinases. Oncotarget 2016;7:70080–70091.

27. Wąsik N, Sokół B, Hołysz M, et al. Serum myelin basic protein as a marker of brain injury in aneurysmal subarachnoid haemorrhage. Acta Neurochir (Wien) 2020;162:545–552.

28. Sakai LY, Keene DR, Renard M, De Backer J. FBN1: The disease-causing gene for Marfan syndrome and other genetic disorders. Gene 2016;591:279–291.

29. Pirruccello JP, Chaffin MD, Fleming SJ, et al. Deep learning enables genetic analysis of the human thoracic aorta. bioRxiv 2020:2020.05.12.091934.

30. Anon. FinnGen Documentation of R4 release. Available at: https://finngen.gitbook.io/documentation/. Accessed December 16, 2020.

31. Jones GT, Tromp G, Kuivaniemi H, et al. Meta-Analysis of Genome-Wide Association Studies for Abdominal Aortic Aneurysm Identifies Four New Disease-Specific Risk Loci. Circ Res 2017;120:341–353.

